# A beneficial synonymous substitution in EF-Tu is contingent on genetic background

**DOI:** 10.1101/2023.09.06.555949

**Authors:** Kaitlyn M. McGrath, Steven J. Russell, Evrim Fer, Eva Garmendia, Ali Hosgel, David A. Baltrus, Betül Kaçar

**Author notes:** To whom correspondence must be addressed.

## Abstract

Synonymous mutations are changes to DNA sequence that occur within translated genes but which do not affect the protein sequence. Although often referred to as silent mutations, evidence suggests that synonymous mutations can affect gene expression, mRNA stability, and even translation efficiency. A collection of both experimental and bioinformatic data has shown that synonymous mutations can impact cell phenotype, yet less is known about the molecular mechanisms and potential of beneficial or adaptive effects of such changes within evolved populations. Here, we report a beneficial synonymous mutation acquired via experimental evolution in an essential gene variant encoding the translation Elongation Factor protein EF-Tu. We demonstrate that this particular synonymous mutation increases EF-Tu mRNA and protein levels, as well as the polysome abundance on global transcripts. Although presence of the synonymous mutation is clearly causative of such changes, we also demonstrate that fitness benefits are highly contingent on other potentiating mutations present within the genetic background in which the mutation arose. Our results underscore the importance of beneficial synonymous mutations, especially those that affect levels of proteins that are key for cellular processes.

**Importance:** This study explores the degree to which synonymous mutations in essential genes can influence adaptation in bacteria. An experimental system whereby an *Escherichia coli* strain harboring an engineered translation protein Elongation Factor-Tu (EF-Tu) was subjected to laboratory evolution. We find that a synonymous mutation acquired on the gene encoding for EF-Tu is conditionally beneficial for bacterial fitness. Our findings provide insight into the importance of the genetic background when a synonymous substitution is favored by natural selection and how such changes have the potential to impact evolution when critical cellular processes are involved.

## Introduction

Synonymous mutations are changes in the codon sequence that do not alter the sequence of the translated peptide. While often considered neutral or silent, evidence suggests that synonymous mutations can affect cellular phenotypes and potentially impact population growth fitness (1–10) as well as gene and protein expression. For example, synonymous mutations may cause modification of mRNA levels by enhancing promoters (8) and altering mRNA folding and/or stability (10–13). Furthermore, changes in mRNA secondary structure can interfere with protein synthesis, disrupting translation speed and accuracy (10, 14–17), which can ultimately impact protein levels. Since translation coincides with protein folding, changes in translation speed and accuracy can influence both protein expression and function (18).

The reported effects associated with codon bias, which refers to the uneven distribution of synonymous codons for the same amino acid in a gene or genome (19), on translation speed and fitness have been mixed. Given the importance of mRNA stability and translation rate for cellular functions, the use of synonymous codons has the potential to impact fitness in a variety of ways. Studies have correlated highly expressed genes with high levels of codon bias which may provide an advantage for cellular fitness (16, 19–23). Conversely, some studies suggest codon bias has minimal or no effect on fitness (24–26), while others show examples of beneficial synonymous mutations with less-preferred codons (6). Taken together, less is known about the molecular mechanisms and potential of beneficial or adaptive fitness effects of such changes within evolved populations.

In this study, we explored the degree in which synonymous mutations influence essential gene adaptation in bacteria building on an experimental system whereby an *Escherichia coli* (*E. coli*) strain harboring an engineered essential gene was subjected to laboratory evolution (27). Specifically, an essential protein in translation, Elongation Factor Tu (EF-Tu), was directly replaced with a phylogenetically inferred ancestor, AnEF, leading to a fitness decrease of the engineered strain (27–28). EF-Tu is an essential GTPase that binds to amino-acylated tRNAs and shuttles them to the A-site of the ribosome **(Fig. 1A)** (29). EF-Tu is critical in mediating the rate and accuracy of translation elongation (30) and is one of the most abundant proteins in the cell, making up ∼6% of the total protein in *E. coli* (31–32).

**Figure 1:**
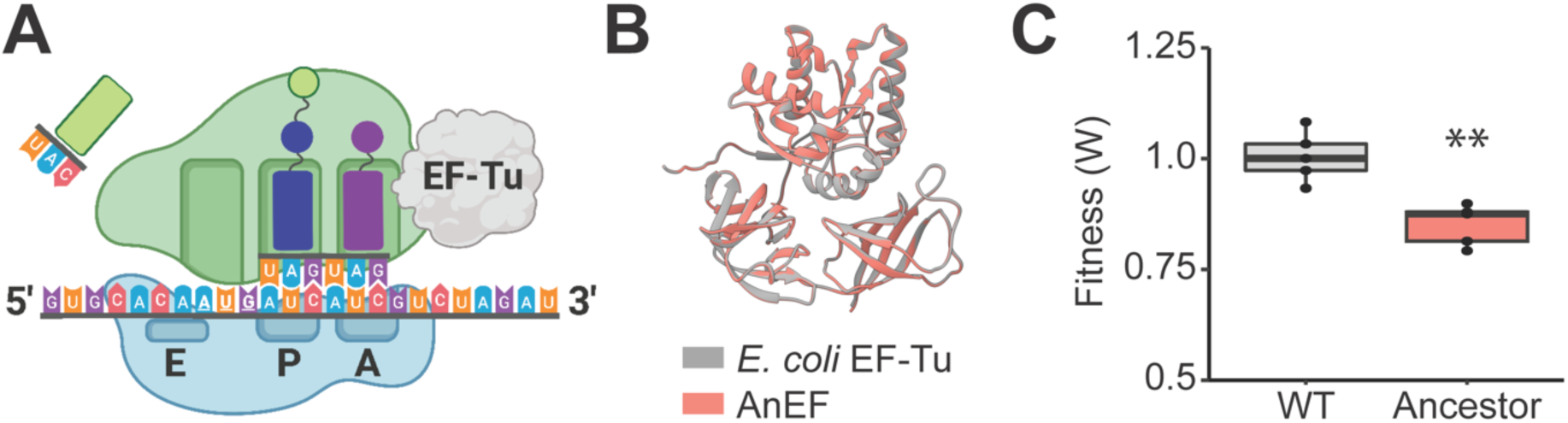
Replacement of EF-Tu with ancestral EF-Tu (AnEF) (A) Overall schematic of interaction between *E. coli* EF-Tu and ribosome (B) Overlay of EF-Tu and AnEF predicted structures results in an RMSD score of 0.162. (C) Competitive fitness of the REL606 harboring AnEF (ancestor) relative to the REL606 Δ*tufA* (wild-type), *t*-test (n = 5).

We report a single synonymous mutation discovered in the *anEF* gene coding region in one of the experimentally evolved engineered *E. coli* populations. Whole genome sequencing reveals that the synonymous mutation in AnEF appears after generation 2000 and sweeps to fixation within the population by the 2500^th^ generation. Genetic and cellular analysis demonstrate that this synonymous mutation is beneficial during laboratory growth and increases the mRNA and protein levels of AnEF, yet these fitness effects are contingent on the presence of one or more additional mutations in the evolved genetic background. Furthermore, we show that this synonymous mutation increases abundance of polysomes during translation. Our results demonstrate that synonymous mutations may play a key role when affecting the levels of proteins that limit key cellular processes.

## Results

In previous work, we engineered and experimentally evolved an *E. coli* strain widely used for laboratory evolution studies, REL606, to replace a single copy of ancestral EF-Tu by substituting the *tufB* gene with an inferred ancestral allele in a strain where the homologous gene copy *tufA* was deleted (*anEF, ΔtufA tufB*::*anEF*) (27). We refer to *ΔtufA tufB*::*anEF* as the ancestor strain hereafter (**Table 1**). *E. coli*’s EF-Tu and AnEF share ∼95% protein sequence identity (**Supp.** Fig 1). Additionally, the predicted structure of AnEF exhibits a Root Mean Square Deviation (RMSD) value of 0.162 when overlaid with *E. coli* EF-Tu (**Fig. 1B**).

**Table 1:** List of *E. coli* strains used in the study. Bacterial strain name and characteristics are indicted. A complete list of strains used for genome engineering can be found under Supplementary Information.

| Strain | Genotype and relevant characteristics |
| --- | --- |
| Ancestor | REL606 $\Delta tufA$, *tufB::anEF* |
| g500 | REL606 $\Delta tufA$, *tufB::anEF* , 500 generations |
| g1000 | REL606 $\Delta tufA$, *tufB::anEF* , evolved 1000 generations |
| g1500 | REL606 $\Delta tufA$, *tufB::anEF* , evolved 1500 generations |
| g2000 | REL606 $\Delta tufA$, *tufB::anEF* , evolved 2000 generations |
| g2500 | REL606 $\Delta tufA$, *tufB::anEF* , evolved 2500 generations |
| g3000 | REL606 $\Delta tufA$, *tufB::anEF* , evolved 3000 generations |
| Evolved (AnEF<sub>C45T</sub>)<br>(isolated clone) | REL606 $\Delta tufA$, *tufB::anEF*, evolved 3000 generations, *anEF* C45T (GTC→GTT) |
| Evolved (AnEF<sub>T45C</sub>)<br>(engineered strain) | REL606 $\Delta tufA$, *tufB::anEF*, evolved 3000 generations, *anEF* T45C (GTT→GTC) |
| Ancestor (AnEF<sub>C45T</sub>)<br>(engineered strain) | REL606 $\Delta tufA$, *tufB::anEF* C45T (GTC→GTT) |

We assessed the effect of AnEF allele replacement into *E. coli* in glucose minimal medium via co-culture competition, of which the fitness was compared to a wild-type strain containing a single copy of the wild-type EF-Tu (*E. coli* REL606 Δ*tufA*). The relative fitness of the modified ancestor strain was ∼0.85 (n = 5, p < 0.01, ANOVA) (**Fig. 1C**). This modified ancestor strain was then subjected to 3000 generations of laboratory evolution through serial propagation of bacterial populations (**Fig. 2A)** (33). An *E. coli* REL606 Δ*tufA* strain was evolved in parallel as a WT control. Using co-culture competition assays, we quantified the fitness of the evolved population relative to the ancestor strain every 1000 generations of the evolution experiment. Generations 1000 and 2000 demonstrate a relative fitness of 1.49 and 1.51, respectively, and generation 3000 displays a relative fitness of generation of 1.65 (n = 5, p < 0.01, ANOVA Tukey HSD) (**Fig. 2B**).

**Figure 2:**
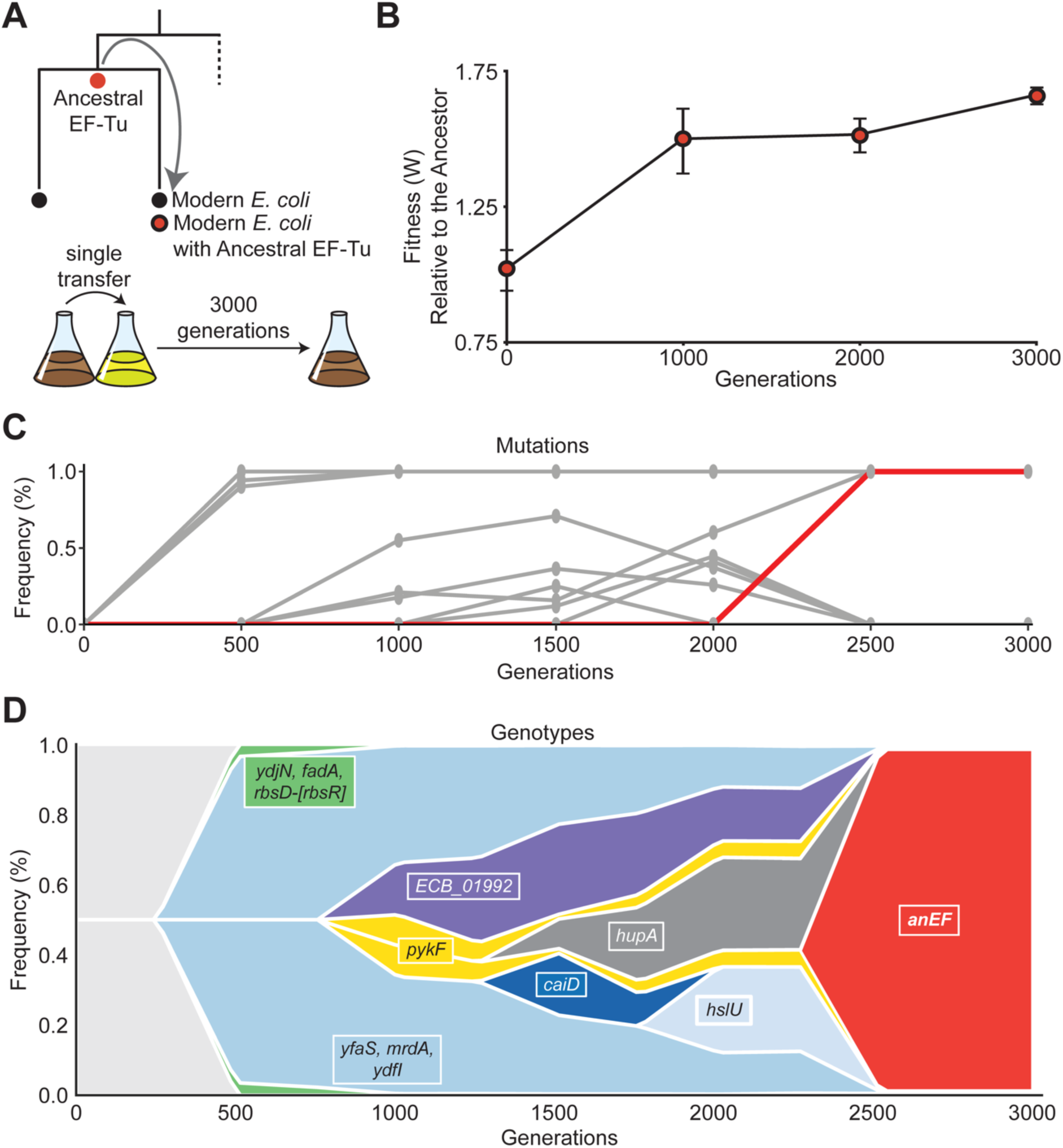
Evolutionary dynamics of an *E. coli* strain harboring an ancestral EF-Tu gene. (A) Engineering and evolution experiment schematics. (B) Change in fitness of evolved population relative to the ancestor. (C) Gene mutation frequencies in evolved population over 3000 generations. *AnEF* synonymous mutation highlighted in red. (D) Muller plot demonstrating the genotype dynamics across laboratory evolution with 3000 generations. Plotted are the mutations in genes that reached a minimum frequency of 25%, and highlighted in red is the selection frequency of the *anEF* gene.

Genomes of evolved populations were sequenced at six different time points: generations 500, 1000, 1500, 2000, 2500, and 3000. Sequencing of the entire mixed population for this experimental lineage revealed 12 mutations that reached more than 25% frequency, however only a handful of mutations became “fixed” (reached a frequency of 100% in the population) in each generation (Fig. 2C**-D, Table S3**). Generation 500 had only 3 fixed nonsynonymous mutations in genes *ydjN* and *fadA*, and a genomic deletion in the *rbs* operon (*rbsD*–[*rbsR*]) (Fig. 2C**-D, Table S3**). Generations 1000 to 2000 acquired 3 additional fixed nonsynonymous mutations in genes *mrdA*, *ydfI*, and *yfaS* (Fig. 2C**, Table S3**). Finally, in generations 2500 and 3000, we identified 2 more fixed mutations, a non-synonymous mutation in gene *pykF* and a synonymous mutation in the replaced gene *anEF* (Fig. 2C**, Table S3**). We next reconstructed the dynamics of substitution associated with each generation time point (Fig. 2D) and identified nonsynonymous mutations that arose independently in four other genes *ECB_01992* (arose in generation 1000), *hupA* (generation 1500), *caiD* (generation 1500), *hslU* (generation 2000) (Fig. 2D). However, these mutations did not reach fixation and instead were eventually lost from the population. The patterns we observed in genotype dynamics is consistent with each detected fixed mutation (Fig. 2C).

### The rise and effect of the AnEF synonymous mutation

Intriguingly at generation 2500, a synonymous mutation appeared in the *anEF* gene coding region. Notably, the synonymous mutation reached 100% fixation rapidly, appearing between generations 2000 and 2500 (Fig. 2C**-D, Table S3**), and ultimately swept the entire population by generation 3000 (Fig. 2D). Specifically, we identified a C > T mutation at nucleotide 45, GT<u>C</u> (Val) → GT<u>T</u> (Val) mutation, in the N-terminal coding region of the *anEF* gene (Fig. 3A, **Table 1**). The presence of the mutation at each generation was successfully verified with whole genome sequencing (**Fig 2C-D, Table S3**).

**Figure 3:**
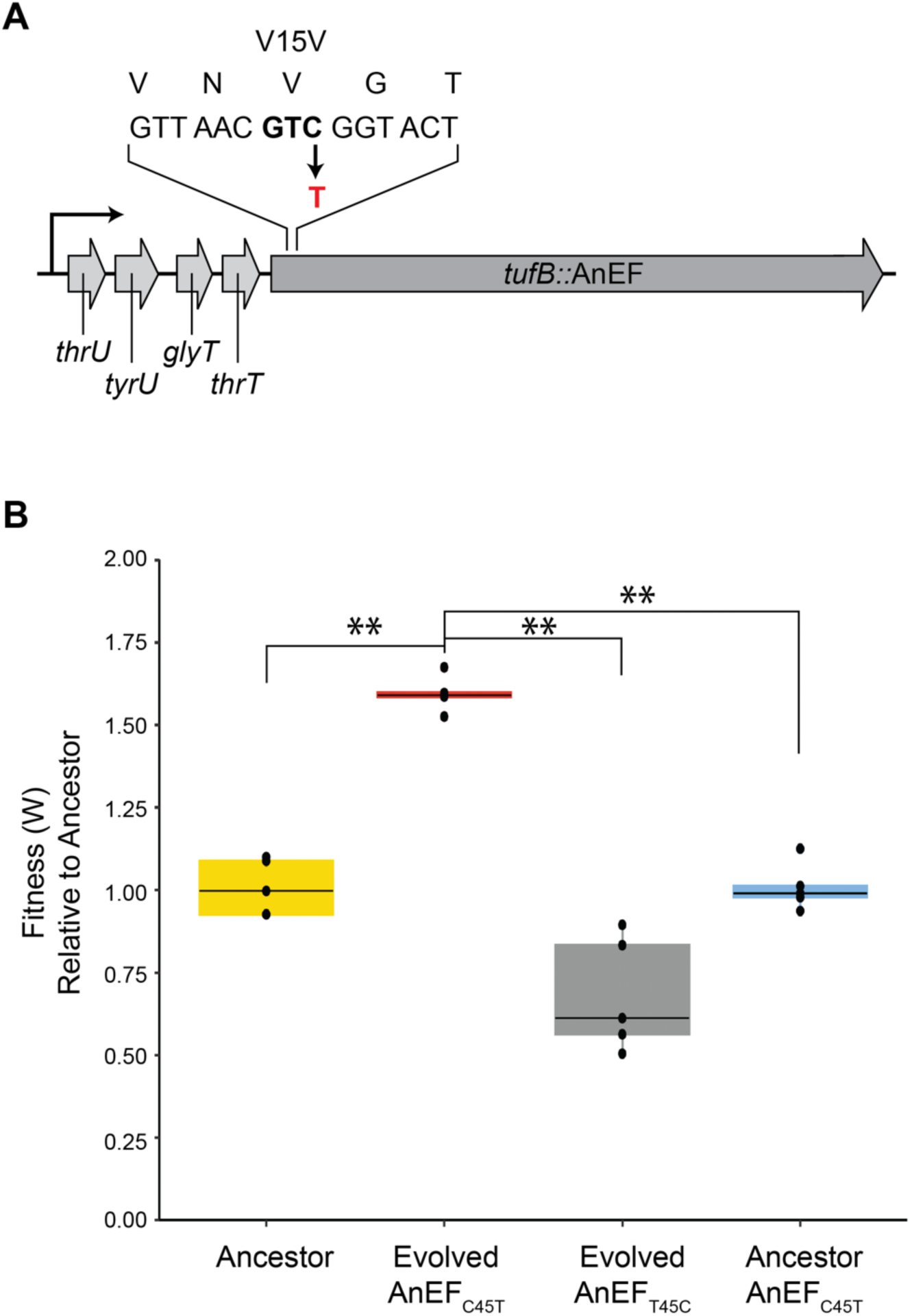
Fitness and growth characteristics of AnEF_C45T_. (A) Schematic of AnEF_C45T_. (B) Fitness phenotype between endogenous and isogenic constructs (n = 5, ANOVA, Tukey’s HSD).

We hypothesized that if the synonymous mutation is beneficial, reverting the evolved mutation in AnEF back to the ancestral nucleotide would negatively affect fitness. To test this hypothesis, we engineered the codon containing the synonymous mutation back to the ancestral codon sequence while keeping the rest of the genetic background (including all other fixed mutations) constant. This new strain is referred to as evolved+AnEF_T45C_ (**Table 1**). Furthermore, to test if the fitness impact of the synonymous mutation was background dependent, we introduced the synonymous mutation (C45T) into the *anEF* unevolved allele in the ancestral background (referred to as ancestor+AnEF_C45T_) (**Table 1**, Methods). All isogenic constructs were confirmed via local DNA sequencing and whole genome sequencing.

### The synonymous mutation is conditionally beneficial depending on the genetic background

We next measured the impact of the C45T synonymous mutation on organismal fitness via co-culture competition assays by calculating relative fitness (W). In agreement with the previously published studies, the replacement of native *E. coli* EF-Tu with the ancestral EF-Tu AnEF causes a 13% decrease in fitness (W=0.87, p < 0.01, ANOVA Tukey’s HSD) (Fig. 1C) (27–28), which upon 3000 generations of laboratory evolution is recovered and increased by 60% relative to the ancestor (Evolved AnEF_C45T,_ W= 1.6, p < 0.01, ANOVA Tukey’s HSD) (Fig. 3B). Substitution of the synonymous mutation with the native codon significantly decreases the fitness of the evolved microbe by 30% (Evolved AnEF_T45C_, W= 0.7, p < 0.01, ANOVA Tukey’s HSD) whereas introducing the synonymous mutation in the ancestral background has no relative fitness benefit (Ancestral AnEF_C45T_, W=0.99, p=0.89, ANOVA Tukey’s HSD) (Fig. 3B). A similar trend is observed when compared relative to REL606 *Δtuf*A. (**Fig. S2**. These results show the strong epistasis between the synonymous mutation and the evolved background, demonstrating that the fitness effect of the synonymous mutation depends on the genetic background.

### AnEF mRNA and protein levels are increased in strains carrying the synonymous mutation

Previous studies have reported that synonymous mutations can impact mRNA and protein levels (6–10). To assess the change in transcript level, we measured mRNA levels via qPCR and calculated ΔΔCq values to compare AnEF mRNA with and without the evolved synonymous mutation (n = 3). Relative to the ancestor, we observed a 3-fold increase in the evolved strain (p < 0.05, *t*-test) and a 2-fold increase in the ancestor with the synonymous mutation (ancestor + AnEF_C45T,_ p < 0.05, *t*-test) (Fig. 4A**, Fig. S3**). The mRNA levels of the evolved strain with the synonymous mutation reverted to the ancestral nucleotide (evolved + AnEF_T45C_) displayed a 30% decrease in AnEF mRNA, however this observation was not significant (p = 0.06, *t*-test) (Fig. 4A**, Fig. S3**). Furthermore, we assessed whether the C45T synonymous mutation has changed the protein levels relative to the ancestor strain (Fig. 4B**, Fig. S3**). AnEF protein levels increase 32.2% in the evolved strain with the C45T synonymous mutation (p < 0.01, *t*-test, Methods) (Fig. 4B**, Fig. S3**). Similarly, the engineered ancestor with the AnEF_C45T_ allele exhibits a 34.2% increase in AnEF protein levels (p < 0.01, *t*-test) (Fig. 4B**, Fig. S3**). Interestingly, reversion of the synonymous mutation back to the ancestral nucleotide in evolved strain (evolved + AnEF_T45C_) leads to a 15.3% decrease in AnEF protein levels (p < 0.01, *t*-test) (Fig. 4B**, Fig. S3**). Taken together, this data suggests that the evolved synonymous mutation in AnEF is correlated with an overall increase in both AnEF mRNA and protein levels.

**Figure 4:**
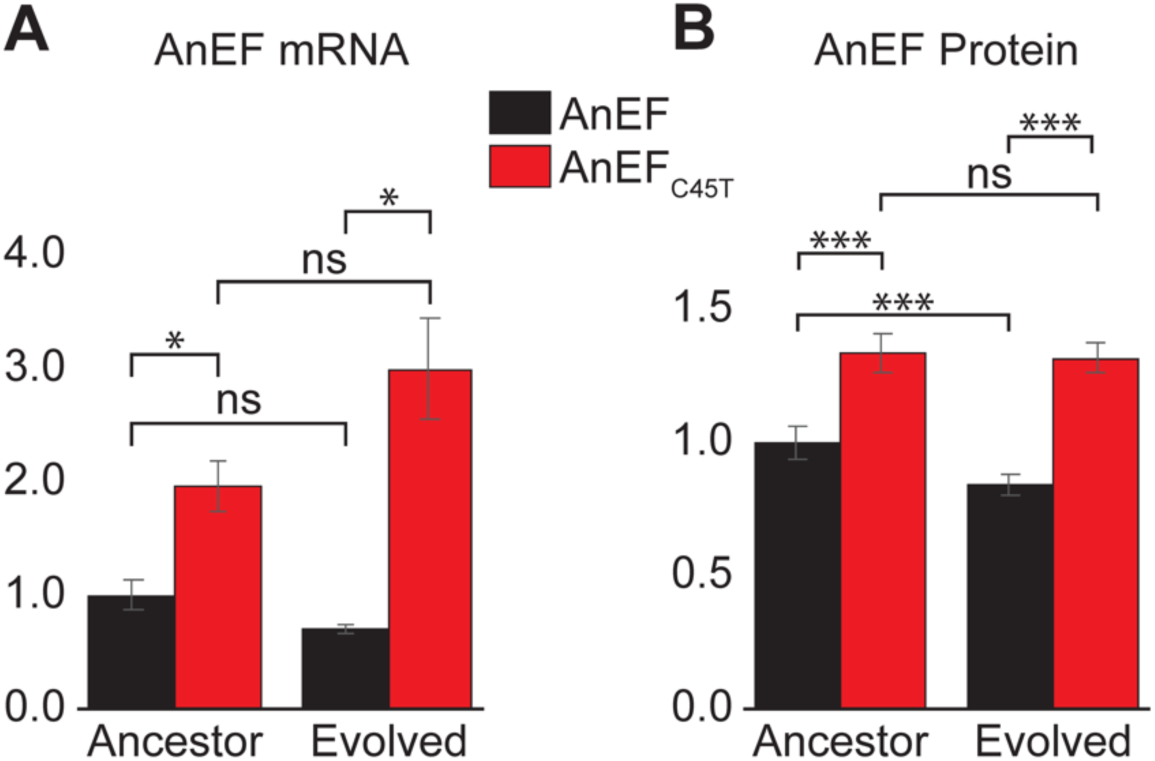
AnEF protein and mRNA levels are increased in strains carrying AnEF_C45T_. (A) qPCR quantification of AnEF mRNA between constructs (n = 3, *t*-test). (B) Western blot quantification of AnEF protein between endogenous and isogenic constructs (n = 3, *t*-test).

### The presence of the synonymous mutation in AnEF leads to an increase in ribosomal abundance in translation elongation

Various studies have demonstrated the link between bacterial growth and ribosomal abundance, showing that growth (and death) rate of bacterial populations is affected by the number of ribosomes (34–36). Elongation Factor proteins are highly expressed in the cell (38) and fulfill a crucial role in the ribosome (39). Thus, upon determining the significant impact the synonymous mutation has on AnEF mRNA and protein levels, we asked if the synonymous mutation affects ribosomal abundance. We hypothesized that the evolved strain with the synonymous mutation would exhibit greater number of ribosomes as it displays an increase in population growth fitness, gene, and protein expression.

We inferred the cellular translatome, which refers to all mRNAs associated with ribosomes in protein synthesis, using polysome profiling to assess the impact the AnEF synonymous mutation has on translation. The translation process is made up of four main steps: initiation, elongation, termination, and recycling, where initiation is the assembly and preparation of the ribosome on a transcript and elongation is the decoding process of protein synthesis (40–41). Polysome profiling provides insight into the translation process by first separating ribosomes via RNA sedimentation and then providing comparative ribosomal abundance for each step of protein synthesis for the corresponding strain. We first assessed the polysome profile of the wild-type REL606 strain. As shown in Figure 5, we used a 10-40% sucrose gradient for sufficient separation of RNA molecules, of which free RNAs (mRNA, tRNA, etc.) are the lightest and therefore will sediment towards the top of the gradient, and the heaviest RNA molecules being the longer stretch of polysomes actively translating along an mRNA transcript. Therefore, the first peak detected on a polysome profiles represents the free RNAs in a cell, followed by peaks associated with the small and large ribosomal subunits, 30S and 50S, respectively. The highest peak shown here represents the assembled 70S monosome, which is a single ribosome on an mRNA transcript, or a ribosome in the initiation phase of translation. Finally, each peak following the monosome represents the stretch of polysomes in the elongation phase of translation (Fig. 5, Methods).

**Figure 5:**
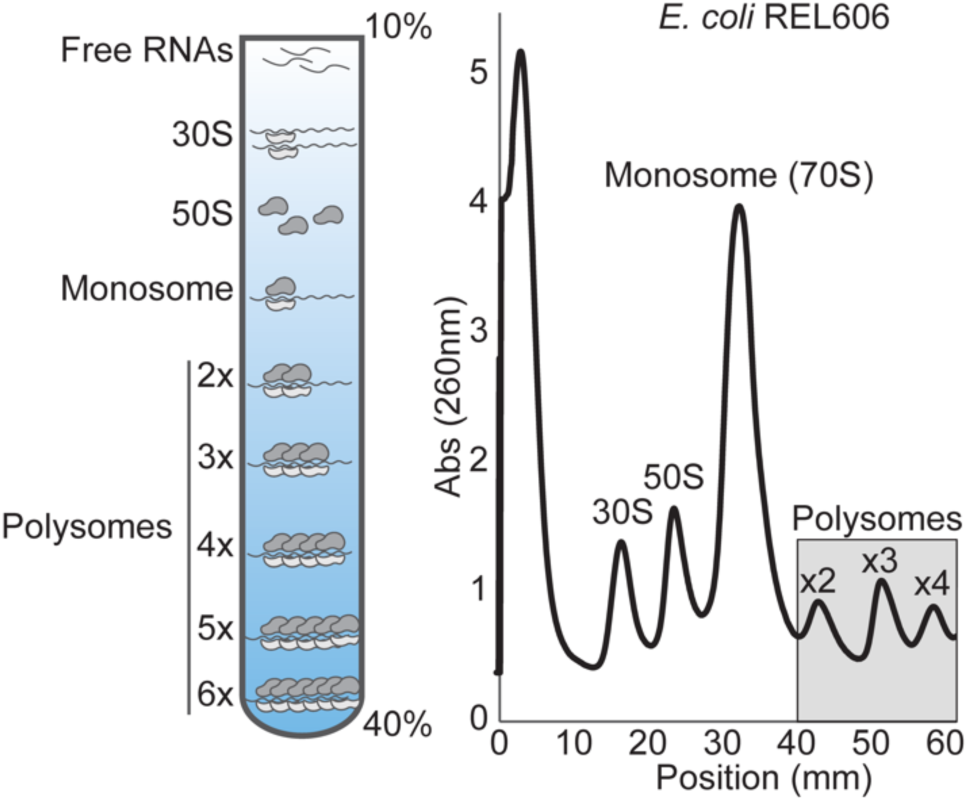
*E. coli* translatome of REL606 assessed via polysome profiling, showing ribosomal abundance measured by 260 nm absorbance along a 10-40% sucrose gradient.

To assess the impact the AnEF synonymous mutation has on translation, we globally quantified the abundance of ribosome footprints by calculating the area under the curve (AUC) for each ribosomal peak and compared relative ribosomal abundance between the endogenous and isogenic strains (n = 6) (Fig. 6, Methods). Relative to the ancestor, the evolved strain has increased levels of the 30S (20%, p < 0.05, two-tailed *t*-test), and 50S (∼13%, p < 0.05, two-tailed *t*-test) ribosomal subunits, as well as an increase in the overall polysome abundance (10%, p < 0.05, one-tailed *t*-test, p ≤ 0.08, two tailed test) (Fig. 6A**, Fig. S4)**. However, compared to ancestor, the evolved strain has decreased levels of 70S monosomes (17%, p<0.05, two-tailed *t*-test) (Fig. 6**, Fig. S4**).

**Figure 6:**
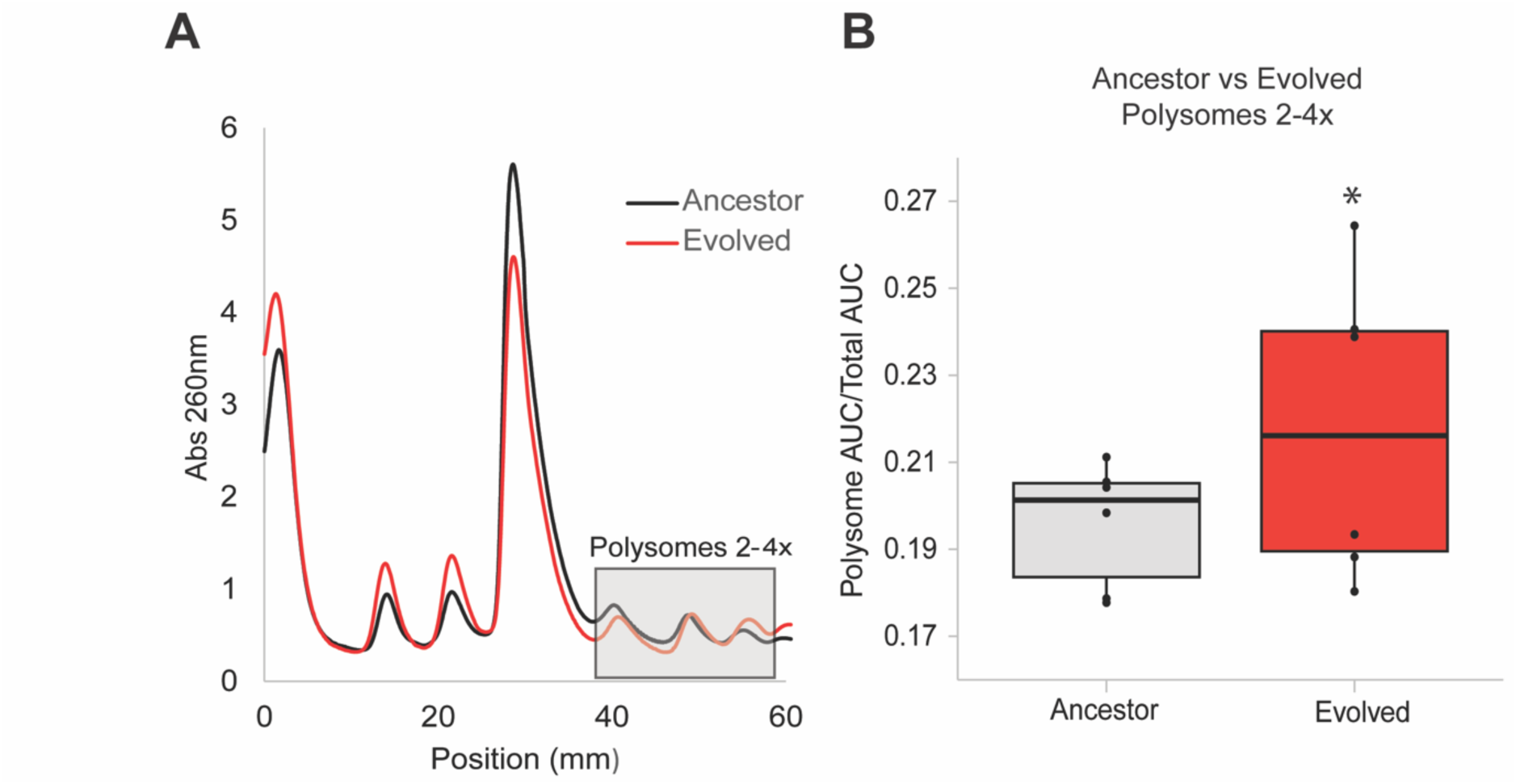
Polysome profiling of the evolved strain (red) relative to the ancestor (black) (B) Quantification of polysome abundance from shaded box in the profiles in generated panel A (n = 6, *t*-test).

## Discussion

Previous studies have shed light on the evolutionary patterns of EF-Tu affecting cellular translation (27–28, 42–44). While translation can be sustained in *E. coli* by the reconstructed ancestral allele of EF-Tu (AnEF), this interchange leads to a decrease in cellular protein synthesis and population fitness (28). Overexpressing the AnEF partially increases *E. coli* fitness (28), demonstrating that fitness may be restricted by the availability of key enzymes that mediate critical cellular processes. We suggest that when protein levels are the limitation for cell growth, synonymous mutations that increase protein levels may lead to beneficial and adaptive outcomes.

The synonymous mutation in AnEF arising during laboratory evolution in *E. coli* is associated with an increase in AnEF mRNA and protein levels (Fig. 4). Intriguingly, a previous study identified that the first 65 codons of *tufB* are important for *tufB* regulation, and that various synonymous mutations near the start codon (including C45) promotes an open conformation of the *tufB* mRNA, which leads to increased EF-Tu expression (45). Since AnEF was inserted in place of *tufB* and shares a nucleotide sequence identity of 87%, it is possible that the observed increase in AnEF mRNA and protein levels can thus be explained by the favored open conformation of AnEF mRNA secondary structure. Moreover, the wild-type *E. coli tufB* codon for valine (position 45) is GTC, which is reported at a frequency of 20% in codon usage bias frequency index (46). Interestingly, the synonymous mutation in AnEF changed the codon from GTC to GTT, which is reported at a higher frequency of 28%, the second most frequent codon used for valine in *E. coli*. To what degree this conversion may be attributable to the codon bias requires further mechanistic investigation.

The synonymous mutation detected in AnEF is also associated with an increase in polysome abundance (Fig. 6**, Fig. S4**). An increase in polysomes represents an increase in the number of elongating ribosomes on global mRNA transcripts. Based on this, we develop three scenarios to interpret our results: (i) The increase in polysomes may reflect the observed increase in AnEF protein levels. The transition of translation initiation to elongation is marked by the first translocation via EF-G (40–41), which is dependent on EF-Tu for bringing a cognate amino-acylated tRNA to form the first dipeptide bond (40–41). Therefore, the boost in AnEF protein abundance may increase the frequency of elongating ribosomes in the cell. (ii) The increase in polysomes may be due to a decline in translation elongation rate. Kaçar et al. (28) shows that protein synthesis rate is reduced when translation is dependent on the EF-Tu ancestral variant. Further, a more ancient EF-Tu variant was demonstrated to have significantly low K_M_ values for dipeptide formation, indicating that ancestral EF-Tus are less efficient at sequestering cognate amino-acylated tRNAs to the ribosome (44). AnEF’s reduced efficiency at sequestering cognate amino-acylated tRNAs to the ribosome could lead to ribosomal stalling in the decoding process. As a result, an increase in stalling may increase the presence in elongating ribosomes. (iii) Finally, a combinatorial effect of the scenarios (i) and (ii) collectively increase cellular translation behavior. The increase in AnEF protein levels equates to an increase in EF-Tu availability for ribosomes transitioning from translation initiation to elongation. However, the boost in AnEF protein amount would not necessarily boost the efficiency of the AnEF function. In sum, the increase in polysome abundance on global mRNA transcripts may be credited to the increase in AnEF protein amount coupled with the AnEF’s kinetic reduction in translation efficiency relative to its modern counterpart.

The AnEF synonymous mutation (AnEF_C45T_) is the cause for the beneficial changes, yet the advantageous impact of the mutation on the population fitness is observed only when the synonymous mutation is present together with the evolved genetic background (Fig. 3). This dependency demonstrates epistatic interactions between the AnEF synonymous mutation and the evolved genetic background, and strongly indicates that there are additional changes required to optimize levels of AnEF in the context of the extant *E. coli* genome. Furthermore, as shown in Fig. 4, the mRNA levels of AnEF_C45T_ are clearly much higher in the evolved background compared to the ancestor, suggesting that epistasis within the evolved background is reinforcing EF-Tu’s role as a limiting factor upon cell growth.

Upon 3000 generations of bacterial evolution, there were 7 additional nonsynonymous mutations that arose and were fixed prior to the synonymous mutation in *anEF* (genes: *mrdA*, *ydfI*, *pykF*, *ydjN*, *yfaS*, *rbsD*-[*rbsR*], and *fadA*) (Fig. 2C**-D & Table S3**). The mutations in *ydjN*, *fadA*, and *rbsD*-[*rbsR*] became fixed in generation 500, where YdjN functions as a L-cysteine transporter (47) and FadA and products of the *rbs* operon function in metabolism (48–49). The genomic deletion in *rbsD*-[*rbsR*] has been witnessed in previous bacterial evolution experiments, in which all twelve parallel evolved populations acquired a loss in the *rbs* operon (involved in ribose catabolism) (50). This *rbs* operon deletion is hypothesized to provide an advantage in glucose minimal medium; thus, we do not suspect it is responsible for the increase in fitness associated with the AnEF synonymous mutation. Mutations fixed in generation 1000 were in genes *mrdA*, *ydfI*, and *yfaS*. The *mrdA* gene (also known as *pbpA*) produces the protein penicillin binding protein 2 (PBP2), which is responsible for maintaining antibiotic sensitivity as well as the rod cell shape in *E. coli* (51); YdfI functions in metabolism (52), while YfaS remains uncharacterized but predicted to be a part of the alpha-2-macroglobulin family (53). The final mutation to fix was in *pykF* in generation 2500 (with *anEF*), the product of which also functions in cell metabolism (54). Changes in some of these genes, for example *mrdA* (55) and *pykF* (56), were previously shown to increase the fitness of REL606 during laboratory evolution. How these background mutations influence AnEF remains to be explored in detail, but their occurrence in such evolution experiments using native, unaltered *E. coli* indicate that phenotypic effects of the synonymous mutation in AnEF are not causally linked on prior mutations in *mrdA*, *pykF*, or the *rbs* operon, and may have arisen due to chance.

Overall, our data underscores the importance of highly expressed proteins in essential cellular processes, as well as the importance of the genetic background when a synonymous substitution is favored by natural selection. The synonymous mutation in AnEF is associated with increased mRNA (Fig. 4A), increased protein (Fig. 4B), and increased polysome abundance (Fig. 6), however, population fitness is only increased when AnEF_C45T_ is coupled with the evolved genetic background (Fig. 3). Taken together, these results demonstrate that synonymous mutations can be beneficial and have the potential to impact evolution when critical cellular processes are involved.

## Acknowledgements

The authors would like to thank Aude Trinquier and Bruno Cuevas-Zuviria for the assistance. This work was supported by a grant from the National Institutes of Health, USA (NIH #T32GM136536) and a NASA Early Career Collaboration Award (KMM), UW Foundation Hiroshi and Sugiyama Fund for Graduate Studies (EF) and the John Templeton Foundation (#61926).

## Author contributions

Conceptualized the study: KM and BK; Performed experiments: KM, SR, EF, EG, AH, BK; Sequence analysis: KM and EF; Wrote the paper: KM and BK; Edited and reviewed: All authors; Final version approved by all authors.

## Materials and Methods

### Media and Culture Conditions

Liquid medium is Luria-Bertani (LB; per liter, 10 g NaCl, 5 g yeast extract, and 10 g tryptone) and Davis minimal medium (25 mg/liter glucose; per liter, 5.34 g K_2_HPO_4_, 2 g KH_2_PO_4_, 1 g SO_4_, 0.5 g sodium citrate, 0.01% MgSO_4_, 0.0002% thiamine (vitamin B1), and 0.0025% glucose). Solid medium is LBA (LB with 1.5% agar) and TA (tetrazolium sugar; per liter, 10 g tryptone, 1 g yeast extract, 5 g NaCl, 1.5% agar, 10 g arabinose, and 0.005% triphenyl tetrazolium chloride (TTC)). All incubations were done at 37°C. LB liquid cultures were shaken at 200 rpm for aeration, and DM25 liquid cultures were shaken at 120 rpm. All media components and chemicals were purchased from Sigma, unless noted otherwise.

### Ancestral protein sequence and structure reconstruction

Ancestral protein AnEF sequence was inferred through ancestral sequence reconstruction of Elongation Factor Tu (EF-Tu) proteins as previously described (57). AnEF structures were predicted by LocalColabFold (58) which is a script to use AlphaFold 2.3.1 on local machines (last used April, 2023). For benchmarking, we also predicted the structure of *E. coli* EF-Tu protein using its protein sequence (P0A6N3.2). The structures were predicted using templates from PDB (*--template*), default number of prediction recycles (*--num-recycle 3*) and amber structure refinement method (*--amber*). The predicted structures were aligned and the RMSD calculation was done using UCSF Chimera MatchMaker (59).

### Strain construction

Ancestral and evolved lineages were derived from *E. coli* B strain REL606 as detailed in (27). The genetic marker TP22-amilCP_opt-kan-sacB-T0 was inserted in intergenic region between *rpoC* and *yjaZ* via dsDNA recombineering (28, 60–61) to link it to the different EF-Tu alleles. The genetic marker and linked EF-Tu allele were moved between strains by P1 *virA* phage mediated transduction (28, 62). Constructs were confirmed via PCR, local Sanger DNA sequencing as well as whole genome sequencing. All sequencing files are deposited to NCBI, Submission ID13305354. A complete list of primers and strain genotypes is listed on Supplementary Tables 1 and 2.

### Experimental evolution

Experimental evolution was carried out in serial dilutions in DM25 liquid medium for 3000 generations (∼6.6 generations per day) as described previously (33) and reported for the AnEF strains (27).

### Whole genome sequencing and analysis

Genomic DNA was extracted from clonal or whole population samples using DNeasy UltraClean Microbial Kit (Qiagen, 12224-50) and shipped to Microbial Genome Sequencing Center (MiGS) for llumina sequencing. All Illumina sequences were analyzed for single nucleotide polymorphisms (SNPs) using the computational pipeline, *breseq* (63). The genomic DNA reference used was REL606 (NCBI RefSeq: NC_012967.1). The C45T synonymous mutation (i.e., V15V) was confirmed with Oxford Nanopore sequencing as well. The quality of Nanopore sequences were checked using fastqc v0.12.1 (64). The reads having base quality lower than 20 were trimmed with Nanofilt v2.8.0 (65) using parameters *-q 20 –headcrop 50*. Trimmed sequences were aligned to the reference genome REL606 (NCBI RefSeq: NC_012967.1) with minimap2 v2.26 (66). The “sam” files that was resulted from alignment were converted into “bam” format and sorted using samtools v1.3.1 (67). The variants were called using NanoCaller v3.0.0 (68). The variants were annotated using vcf-annotator 0.7 (69).

### Bacterial growth curves and doubling time calculations

Strains were grown in LB medium. Each strain was diluted into 10 technical replicates at a dilution of 0.01 at OD_600_, grown at 37℃, and shaken at 200 RPM. Data was collected every 15 minutes at OD_600_ for 24 hours using a BMG CLARIOstar plate reader. Growth curve experiments were run alongside an LB blank control and all values were blank-corrected. Subsequently, all values were calculated in Rstudio using growthcurver and plotted using ggplot2, as provided in the GitHub https://github.com/kacarlab/EFTUSyn.

### RNA extractions, cDNA synthesis, and quantitative PCR

Strains were grown in LB medium and collected at 0.3-0.6 OD_600_ for RNA extraction. Pellets were lysed and prepared following the standard protocol in Rneasy kit (Qiagen, Cat#). Following RNA extraction, samples were removed of any contaminating genomic DNA using the Dnase I standard protocol (Invitrogen, 18068015). Following Dnase treatment, RNA extracts were synthesized into cDNA using reverse transcriptase SuperScript™ IV First-Strand Synthesis System (Invitrogen, Cat#). Finally, all cDNA samples were run on PCR to confirm no genomic DNA contamination before quantitative PCR. All cDNA samples (3 biological replicates) were run in SsoAdvanced™ Universal SYBR® Green Supermix (Bio-Rad, 1725271) in 3 technical replicates for 40 cycles. All primers are listed on **Supplementary Table 2**. ΔΔCq values were calculated using housekeeping gene *rpoB* as a control.

### Immunoblotting experiments

Strains were grown in LB rich medium and collected at 0.3-0.6 OD_600_ for cell lysis (3 biological replicates). Cell pellets were lysed in 300 µL of lysis buffer (10 mL BugBuster^®^ [Millipore 70584-4], ¼ tablet cOmplete™ Protease Inhibitor Cocktail EDTA-Free [Roche]) for 20 min at room temperature and centrifuged at 13,000 rpm at 4℃ for 30 min to clear all cell debris. Protein concentrations were measured using BCA assay (Thermofisher 23227) and stored at −20℃. Whole cell lysates were linearized at 95°C for 5 minutes and were run on a 12% resolving Tris-Glycine SDS-PAGE gel at a constant voltage of 125 V. Protein bands were then transferred to a nitrocellulose membrane at a constant amperage of 400 mA for 45 min. All membranes were blocked with 5% nonfat milk for 1 hour at room temperature. To visualize EF-Tu bands, membranes were probed with 1:1,000 EF-Tu monoclonal antibody (Hycult Biotech mAb 900; Cat#: HM6010) for at 4℃ overnight, followed by incubation with 1:10,000 IRDye® 680RD anti-mouse secondary antibody (LI-COR, 929-70050) for 1 hour at room temperature. All membranes were imaged using the LI-COR Odyssey® XF. To normalize for band quantitation, membranes were stained with GelCode Blue Stain (Thermofisher, 24590) for 5 min and destained for 10 minutes using a destaining solution composed of 40% dH_2_0, 10% acetic acid, and 50% methanol. Bands were quantified using ImageJ and statistics were calculated using Rstudio.

### Fitness assays

All competition assays were followed per published Ara-/+ competition protocol (33). Mixed populations were revived with 100 μL from a glycerol stock into 10 mL of LB medium and grown overnight shaking at 250 RPM at 37 ℃. Clonal populations were revived with 2 μL from a glycerol stock. The next day, each mixed population or clonal strain was preconditioned by diluting 1:10,000 in 10 mL of DM25 medium (5 replicates each) and grown overnight shaking at 250 RPM at 37 ℃. To begin the competition, 2 selected strains were combined in equal amounts (50 μL) into 10 mL of DM25, plated 100x diluted on TA agar (*d_0_*) and grown for 24 hours shaking at 250 RPM at 37 ℃. After 24 hours, the competition assay was plated 100x diluted on TA agar a second time (*d_1_*) and grown at 37 ℃ overnight. All plates were imaged after 24 hours. To calculate relative fitness (W), we calculated the ratio of each strain’s Malthusian parameters (*M_A_* and *M_B_*).

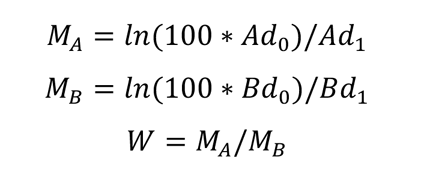

### Polysome profiling

Samples were prepped in accordance with Qin & Frederick (70). Each strain was grown to mid-log (OD_600_ 0.3-0.6) at 37℃ and shaking at 200 rpm. At mid-log, 35 mL of culture was collected, pelleted at 4,000 rpm at 4℃ for 5 minutes, resuspended in 500 µL of chilled lysis buffer (10 mM Tris-HCl pH 8.0, 10 mM MgCl_2_, 1 mg/mL Lysozyme), and flash frozen in liquid nitrogen. The lysates were then thawed in ice water and immediately refrozen in liquid nitrogen and stored at −80℃. To separate polysomes, lysates were thawed on ice and resuspended with 15 µL of 10% sodium deoxycholate. The lysates were then cleared of cell debris at 10,000 rpm at 4℃ for 10 minutes. A normalized volume of 500,000 ng of RNA was carefully loaded on top a sucrose gradient (10-40% sucrose, 20 mM Tris HCl pH 8.0, 10 mM MgCl_2_, 100 mM NH_4_Cl, 2 mM DTT, assembled using Biocomp Gradient Master™) and ultracentrifuged (Beckman Coulter Optima™ XE-90, Rotor SW41Ti) for 3 hours at 35,000 rpm at 4℃. To collect polysome profiles, samples were collected, and RNA was measured at 260 nm using the Biocomp scanner, the overall assay was repeated twice. Ribosome abundance was measured by calculating the area under the curve (AUC) for peaks corresponding to the 30S subunit, 50S subunit, 70S monosome, and polysomes comprising 2-4 ribosomes. Subsequently, peak areas were normalized by the total area under the curve (30S + 50S + 70S + polysomes). AUCs were generated by averaging across both trials.

### Statistical tests

Data analysis was done using Rstudio, ImageJ, and Excel. All data replicates were tested for statistical significance using paired, two-tailed t-tests and ANOVA unless stated otherwise.

## Data availability statement

All raw data are available on GitHub (https://github.com/kacarlab/EFTUSyn). All sequences are available on NCBI Database SUB13308871.

## Notes

### Competing Interest Statement

The authors have declared no competing interest.

