## Supplemental Information for "A beneficial synonymous substitution in EF-Tu is contingent on genetic background"

**Supplementary Table 1**: Strains and genetic markers used in this study

| **Strain** | **Species, genotype, or characteristics** |
|---|---|
| REL606 (WT) | *Escherichia coli*, F-, tsx-467(Am), araA230, lon-, rpsL227(strR), hsdR-, [mal+](LamS) |
| REL607 | REL606 *araA* D92G (GAC→GGC) |
| Δ*tufA* | REL606 Δ*tufA* |
| Ancestor | REL606 Δ*tufA*, *tufB*::*AnEF* |
| Evolved | REL606 Δ*tufA*, *tufB*::*AnEF* V15V (GTC→GTT), *mdrA* T461P (ACC→CCC), *ydfI* Insertion +9 bp, *pykF* T278P (ACC→CCC), *ydjN* Δ24 bp, *yfaS* R900C (CGT→TGT) |
| CH3998 | MG1655 yfaH::[TP22-amilCP_opt-kan-sacB-T0] |
| CH1940 | MG1655 / pSIM5-tet |
| CH6523 | REL606 Δ*tufA, tufB*::*AnEF,* TP22-amilCP_opt-kan-sacB-T0 in intergenic region between *rpoC* and *yjaZ* |
| CH6557 | REL606 Δ*tufA, tufB*::*AnEF*$_{C45T}$, evolved for 3000 generations, TP22-amilCP_opt-kan-sacB-T0 in intergenic region between *rpoC* and *yjaZ* |
| CH6556 | REL606 Δ*tufA, tufB*::*AnEF*, rest of genome evolved for 3000 generations. TP22-amilCP_opt-kan-sacB-T0 in intergenic region between *rpoC* and *yjaZ* |
| CH6585 | REL606 Δ*tufA, tufB*::*AnEF*$_{C45T}$, rest of genome ancestral. TP22-amilCP_opt-kan-sacB-T0 in intergenic region between *rpoC* and *yjaZ* |
| g500 | REL606 Δ*tufA*, *tufB*::*AnEF*, evolved 500 generations (MB+2 500) |
| g1000 | REL606 Δ*tufA*, *tufB*::*AnEF* , evolved 1000 generations (MB+2 1000) |
| g1500 | REL606 Δ*tufA*, *tufB*::*AnEF* , evolved 1500 generations (MB+2 1500) |
| g2000 | REL606 Δ*tufA*, *tufB*::*AnEF* , evolved 2000 generations (MB+2 2000) |
| g2500 | REL606 Δ*tufA*, *tufB*::*AnEF* , evolved 2500 generations (MB+2 2500) |
| g3000 | REL606 Δ*tufA*, *tufB*::*AnEF* , evolved 3000 generations (MB+2 3000) |
| Evolved (AnEF$_{C45T}$) | REL606 Δ*tufA*, *tufB*::*AnEF*, evolved 3000 generations, *AnEF* C45T (GTC→GTT) |
| Evolved (AnEF$_{T45C}$) | REL606 Δ*tufA*, *tufB*::*AnEF*, evolved 3000 generations, *AnEF* T45C *(*GTT→GTC) |
| Ancestor (AnEF$_{C45T}$) | REL606 Δ*tufA*, *tufB*::*AnEF* C45T (GTC→GTT) |

**Supplementary Table 2**: Oligonucleotide primers used in this study

| **Oligo name** | **Sequence (5' to 3')** |
|---|---|
| 606_ksacBblue_linketufB_fw | AGGGAAAGAGCATTTGTCAGAATATTTAAGGAATTTCTGAATCAAAGGGAAAACTGTCCA |
| 606_ksacBblue_linketufB_rv | GTTTCAGGATATTAGTCATCTCTACATTGATTATGAGTATAAAATGAGACGTTGATCGGC |
| 606_rpoC_out_fw | GTTATCGTGGGTCGTCTG |
| 606_yjaZ_out_rv | TGTTTTTCTTCGTTCGTCTG |
| kan_out | GTCATAGCCGAATAGCCTCTCCAC |
| sacB_out | GCTGTACCTCAAGCGAAAGG |
| tufB_coli_fw | TTCTTTTCTCCTCCCTGT |
| tufB_coli_rv | GGCAAACCAAATCGAAAC |
| qPCR_tufB_fwd | AACCGCACGTTAACGTCGGT |
| qPCR_tufB_rev | GCCAGTACGGTAGTGATTGCAGC |
| qPCR_rpoB_fwd | TAAGGTAACGCCGAAAGGTG |
| qPCR_rpoB_rev | CAGAGGCTTTCTCACCGAAG |

**Supplementary Table 3** – Distribution of fixed mutations across 3000 generations in experimentally evolved *E. coli* REL606 (*tufA*Δ, *tufB*::*AnEF*).

| gene | mutation | g0 | g500 | g1000 | g1500 | g2000 | g2500 | g3000 |
|---|---|---|---|---|---|---|---|---|
| *AnEF* | V15V (GTC→GTT) | 0% | 0% | 0% | 0% | 0% | 100% | 100% |
| *mrdA* | T461P (ACC→CCC) | 0% | 90.2% | 100% | 100% | 100% | 100% | 100% |
| *ydfI* | Insertion +9 bp | 0% | 94.2% | 100% | 100% | 100% | 100% | 100% |
| *pykF* | T278P (ACC→CCC) | 0% | 0% | 21% | 15.8% | 60.2% | 100% | 100% |
| *ydjN* | Δ24 bp | 0% | 100% | 100% | 100% | 100% | 100% | 100% |
| *yfaS* | R900C (CGT→TGT) | 0% | 94.3% | 100% | 100% | 100% | 100% | 100% |
| *rbsD*–[*rbsR*] | Δ5,379 bp | 0% | 100% | 100% | 100% | 100% | 100% | 100% |
| *fadA* | Insertion +3 bp | 0% | 100% | 100% | 100% | 100% | 100% | 100% |

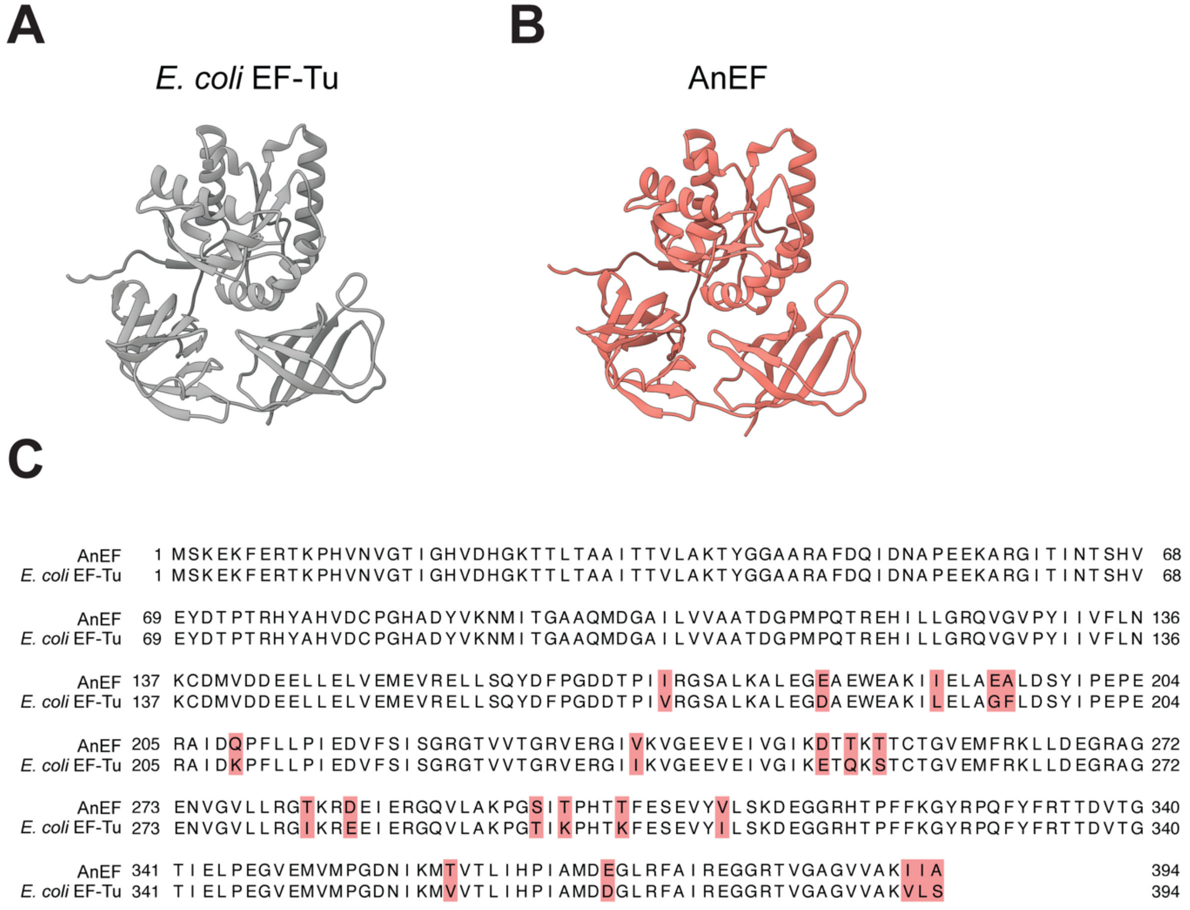


**Supplementary Figure 1**: Comparison of *E. coli* EF-Tu (A) and AnEF (B) protein structures and sequences (C). Predicted structures generated using AlphaFold (see Methods). Alignment of *E. coli* EF-Tu and AnEF protein sequences using MAFFT, alignment algorithm L-INS-i. Differences in sequence highlighted in red. Protein sequence identity ~95%.

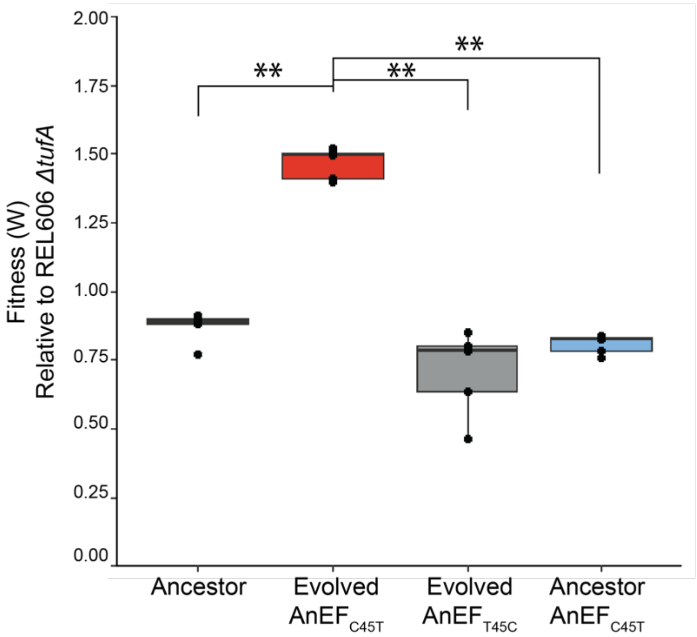


**Supplementary Figure 2**: Fitness characteristics of isogenic constructs relative to REL606 *Δtuf*A (n = 5, ANOVA, Tukey's HSD)

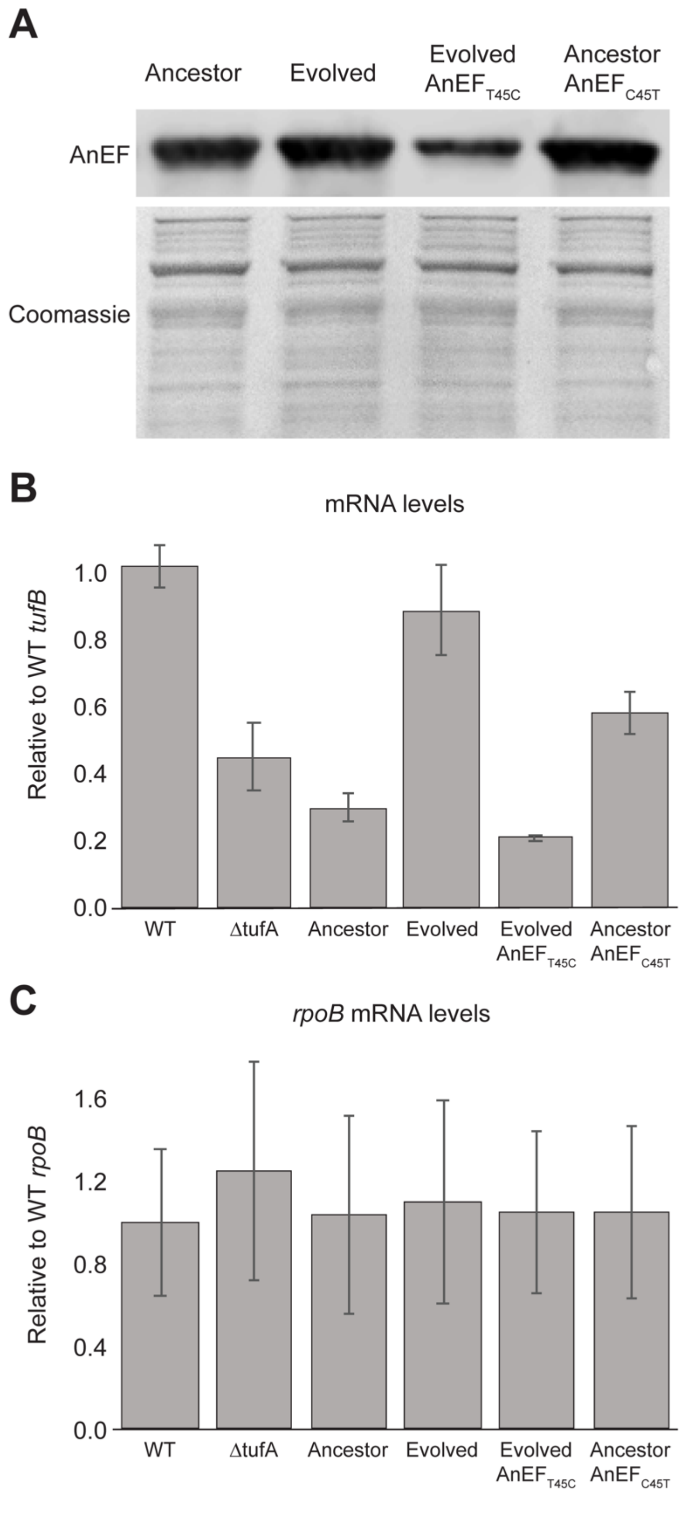


**Supplementary Figure 3**: AnEF protein and RNA levels. (A) AnEF protein levels represented via immunoblot. Total protein was stained using Coomassie. (n=9) (B) qPCR quantification of *AnEF* and *tufB* mRNA, including REL606 (WT) and REL606 ΔtufA. (n=3) (C) qPCR quantification of control *rpoB* mRNA from (B).

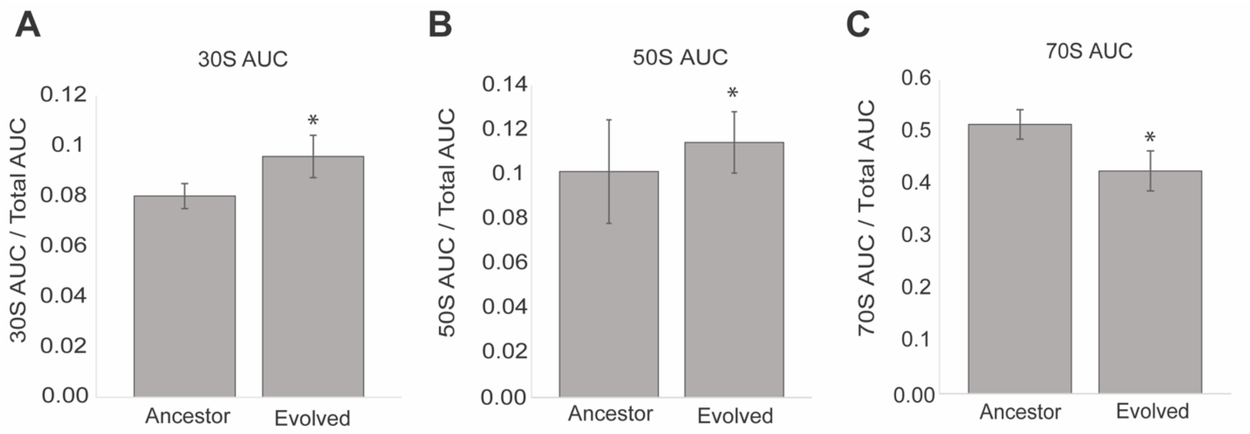


**Supplementary Figure 4**: Polysome profile average area under the curve (AUC) quantification. Comparing the following ribosomal components in ancestor vs. evolved strains: (A) 30S ribosomal subunits, (B) 50S ribosomal subunit, (C) 70S monosomes.
